# Exploring the Upper Boundaries of Distance-Based Protein–Ligand Binding Affinity Prediction: Accuracy Improvements via Linear Regression and Artificial Neural Networks

**DOI:** 10.1101/2025.10.07.680851

**Authors:** Yong Xiao Yang, Bao Ting Zhu

## Abstract

The traditional scoring functions for protein–ligand binding affinity prediction usually adopt fixed mathematical formulas to characterize atomic interaction energies, where electrostatic interactions are described by the 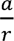 function and van der Waals interactions by the 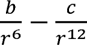 function. These fixed power exponents of distance-based variables severely limit the predictive flexibility and generalization of conventional scoring models. To address this limitation, this study proposes an improved prediction strategy with unrestricted power exponents for distance-dependent atomic interaction variables. To explore the upper boundaries of the predictive powers, the optimized distance-based descriptors were integrated with linear regression and artificial neural network algorithms to construct novel models for protein–ligand binding affinity and binding energy prediction. All linear, artificial neural network, and hybrid models in this work were trained and validated on the latest high-quality dataset from PDBbind (version 2024). Quantitative evaluation based on the Pearson’s correlation coefficient (*R*) demonstrated that the optimized linear models achieved a moderate predictive performance with *R* values ranging from 0.700 to 0.800 on the high-quality affinity dataset, while the artificial neural network-based hybrid models yielded substantially improved accuracy, with *R* values between 0.800 and 0.900. Comparative analysis verified that the optimal models established in this study outperform previously-reported state-of-the-art linear and machine learning-based scoring functions. Specially, this work indicates that the scoring functions with explicit mathematical expressions can achieve a high accuracy close to the experimental error for affinity prediction in a dataset composed of several thousand protein–ligand complexes. The overall results indicate that rationally-optimized power exponents for distance-based variables can effectively enhance the predictive accuracy of protein–ligand binding affinity prediction, providing a reliable and efficient tool for computational drug discovery.

**GRAPHICAL ABSTRACT:** 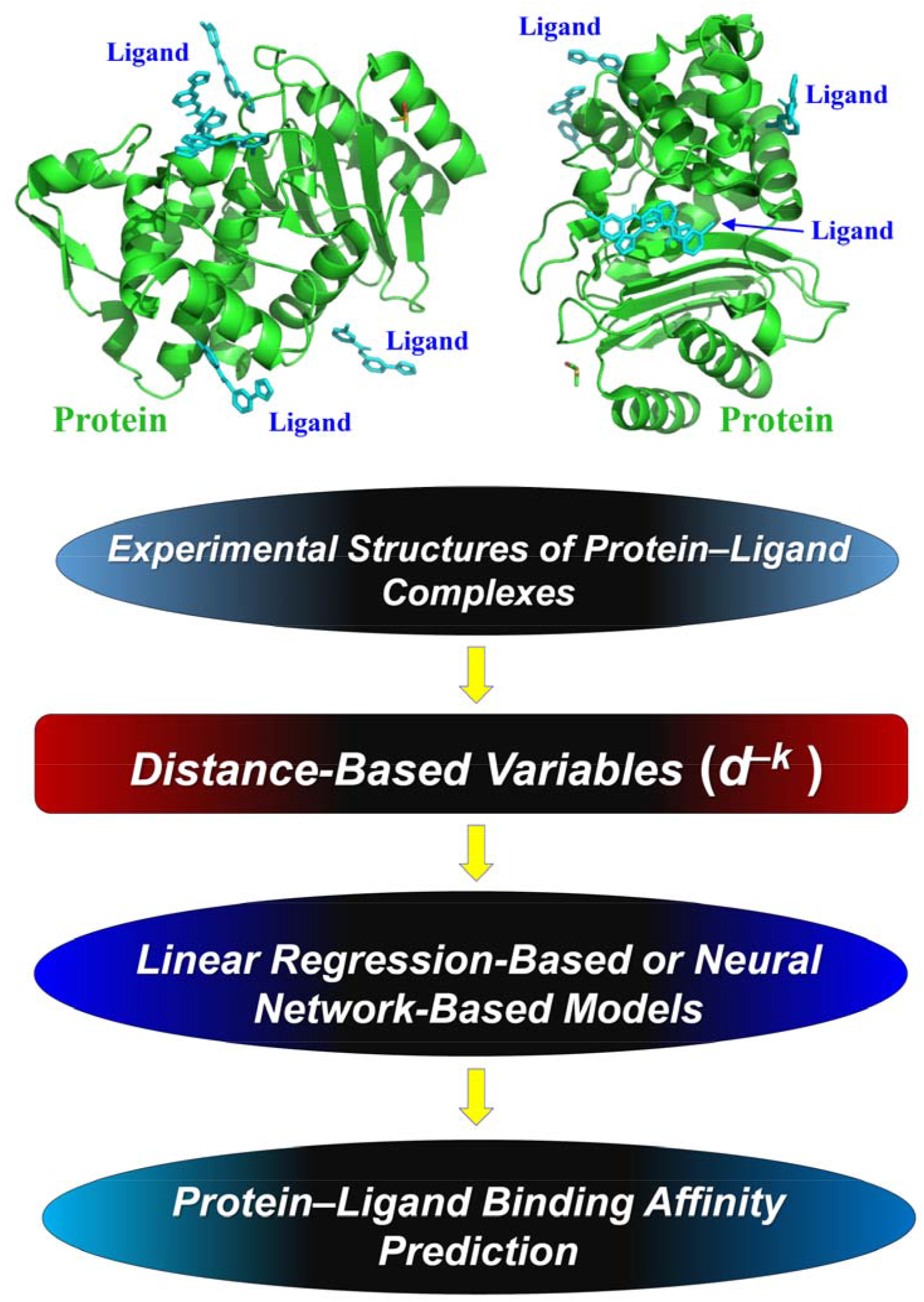

**HIGHLIGHTS:**

- Linear, artificial neural network and hybrid models were constructed using the latest PDBbind v.2024 dataset, with systematic optimization of distance-dependent energy descriptors to explore the upper boundaries of the predictive powers of distance-based variables in protein–ligand binding affinity prediction.
- Relaxing fixed power exponents of distance terms in classic interaction functions achieves finer atom-level depiction of distance-energy relationships in intermolecular interactions.
- The proposed models exhibit superior predictive performance. Specially, this work indicates that the scoring functions with explicit mathematical expressions can achieve a high accuracy close to the experimental errors for affinity prediction in a dataset composed of several thousand protein–ligand complexes.
- This work can be viewed as a new starting point for extending and improving the performances of the traditional energy terms-based scoring functions for binding affinity prediction.

## INTRODUCTION

Protein–ligand binding affinity quantifies the binding strength between ligand molecules (*e.g.*, drug candidates) and target receptor proteins. As a fundamental biophysical parameter, protein–ligand binding affinity not only characterizes the intrinsic interaction patterns of protein–ligand complexes, but also serves as a key indicator guiding rational drug design and lead compound optimization (Mozaffari et al., 2025). Despite its importance and usefulness, experimental determination of the protein–ligand binding affinity remains inherently labor-intensive, time-consuming, and costly, which greatly limits its practical use in high-throughput screening and iterative optimization of drug molecules. To overcome these experimental bottlenecks, computational prediction methods have emerged as a cost-effective and efficient alternative, which not only enables rapid evaluation of protein–ligand binding properties, but also substantially accelerates the drug development pipeline (Dong et al., 2025; Su et al., 2019).

At present, the mainstream computational approaches for predicting protein–ligand binding affinity and binding energy can be broadly classified into two main categories: the molecular dynamics (MD) simulation-based methods and the empirical scoring function-based methods (Bao et al., 2020). While MD simulations deliver rigorous and physically accurate binding predictions by capturing dynamic molecular interactions, their practical applications are severely constrained by excessive computational resource consumption and long calculation cycles, making them practically unsuitable for large-scale dataset screening. By contrast, scoring function-based approaches compromise partial physical rigor for superior computational efficiency, which exhibit unique advantages in high-throughput affinity prediction for massive protein–ligand complex datasets. Traditional scoring functions such as X-Score (Wang et al., 2002), APBScore (Bao et al., 2020), Lin_F9 (Yang and Zhang, 2021), AA-Score (Pan et al., 2022), and LeScore (Xie et al., 2025) are typically parameterized and trained on experimentally-verified structural and affinity data. However, most conventional scoring functions with fixed explicit mathematical formulas suffer from limited prediction generalization and robustness, generally underperforming advanced machine learning-based prediction models that adopt flexible implicit fitting strategies (Lu et al., 2019; Su et al., 2019; Wang and Zhang, 2017; Wang et al., 2002; Yang and Zhang, 2022) and the deep learning-based models with more parameters and complexity (Gao et al., 2026; Gu et al., 2023; Samudrala et al., 2025; Wang et al., 2024; Wang, 2024; Yang et al., 2025; Zhang et al., 2026).

Specifically, the following earlier research works have prompted or inspired our present study, with a goal to make practical improvements in certain aspects:

□ While mathematical equations describing electrostatic and van der Waals interactions in the traditional potential energy function (Levitt, 2001) facilitated the calculation of the binding energy in molecular interactions, these equations, in some ways, also restricted the computational accuracy for binding energy or affinity prediction. Therefore, the present work seeks to extend the mathematical expression of the traditional energy terms.
□ Past linear scoring functions have adjusted the weights and contributions of the electrostatic, van der Waals and hydrogen bonding interaction energy terms and have also accounted for the deformation penalty and hydrophobic effect in protein–ligand complex (Bao et al., 2020; Wang et al., 2002). However, the performances of the traditional linear scoring functions in the CASF-2016 Core Set have not exceeded 0.700 in terms of *R* values. As such, the present study seeks to jointly adjust all coefficients and weights of the original distance-based variables to improve the *R* values.
□ The existing machine learning-based scoring functions (Wang and Zhang, 2017; Yang and Zhang, 2022) integrated different geometric and energy terms in nonlinear ways without explicit mathematical expressions. The performances were further improved to 0.845 in terms of *R* values in the CASF-2016 Core Set. This study seeks to explore the upper boundaries of the predictive powers of the original distance-based variables using artificial neural network.

Therefore, the present research aims to explore and optimize distance-based protein–ligand binding affinity prediction models by integrating linear regression and artificial neural network algorithms to detect the upper boundary of the predictive powers of distance-based variables. Following the variable expansion strategy reported in previous studies (Yang and Zhu, 2025), we refine the power exponent configuration of distance-dependent interaction variables in traditional empirical energy terms. Specifically, electrostatic interaction energy is described via the classical 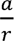 mathematical formulation, while van der Waals interaction energy is characterized by the standard 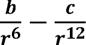 potential function, where ***r*** represents the distance of interacting atom pairs, and ***a***, ***b***, and ***c*** denote tunable model parameters. To further improve the reliability and generalization capability of the constructed models, we adopt the latest and high-quality experimental structural dataset from PDBbind (version 2024) which was compiled based on the available data in the Protein Data Bank at the beginning of 2024 (Berman et al., 2000; Liu et al., 2017; Wang et al., 2004) for model training and validation. Systematic performance validation demonstrates that the optimized models proposed in this work, in some instances, achieve superior prediction accuracy compared with previously developed state-of-the-art sophisticated prediction methods.

## RESULTS

The overall workflow of this study is illustrated in **Fig. 1**. The methodological procedures are summarized as follows: ***i*.** data filtering from the PDBbind v.2024 dataset (Wang et al., 2004); ***ii*.** atom type assignment for experimental complex structures using Open Babel 2.4.1 (O’Boyle et al., 2011); ***iii*.** calculation of distance-based structural descriptors; ***iv*.** development of pre-linear and neural network models using the derived distance-based descriptors; ***v*.** generation of linear models by integrating the predicted values of the selected pre-linear models and creation of hybrid models by integrating the predicted values of the selected neural network models; ***vi*.** development of optimized final hybrid models with variable power exponents on the basis of the preliminary hybrid models. All the scoring functions or models developed in this work are empirical, rather than theoretically derived. One single fixed power exponent exists in each of the distance-based descriptors, pre-linear models, neural network, and hybrid models, and multiple power exponents exist in the linear and final hybrid models. The details of the model generation process are described in **Materials and Methods** section.

**Figure 1.**
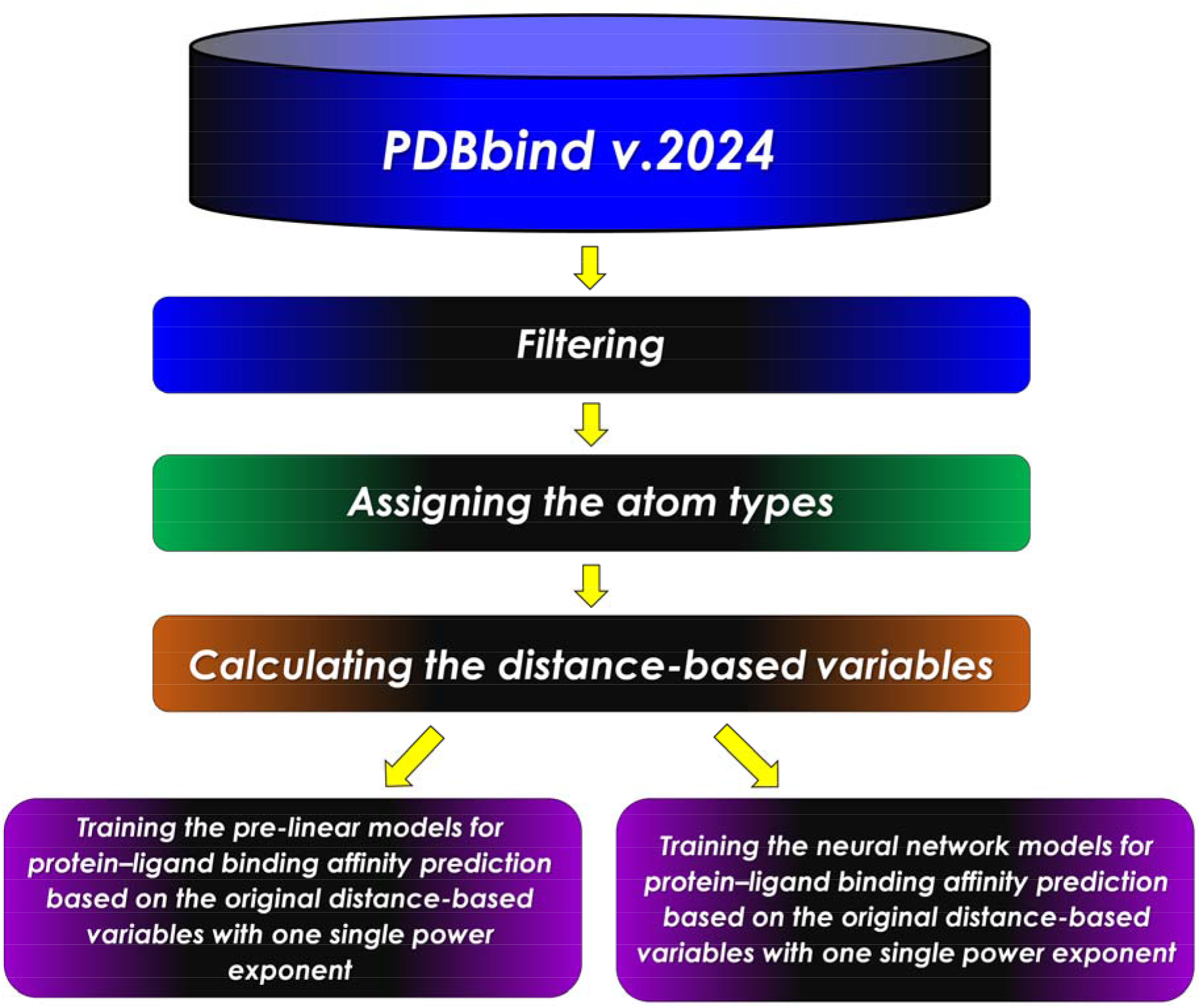
Flowchart of this work for distance-based protein−ligand binding affinity prediction. The steps include: **1**) filtering the data; **2**) assigning the atom types; **3**) calculating the distance-based variables; **4**) training pre-linear and neural network models; **5**) training the linear models based on the predicted values of the selected pre-linear models and training the hybrid models based on the predicted values of the selected neural network models; **6**) training the final hybrid models based on the predicted values of the selected hybrid models.

A total of 13,733 protein–ligand complexes were included in the dataset. Approximately 40% of these complexes contain multiple identical ligand molecules bound to distinct sites within the same protein structure (**Fig. 2A, 2B**). For most multi-ligand complexes, the number of identical bound ligand molecules per protein structure is no more than five (**Fig. 2B**).

**Figure 2.**
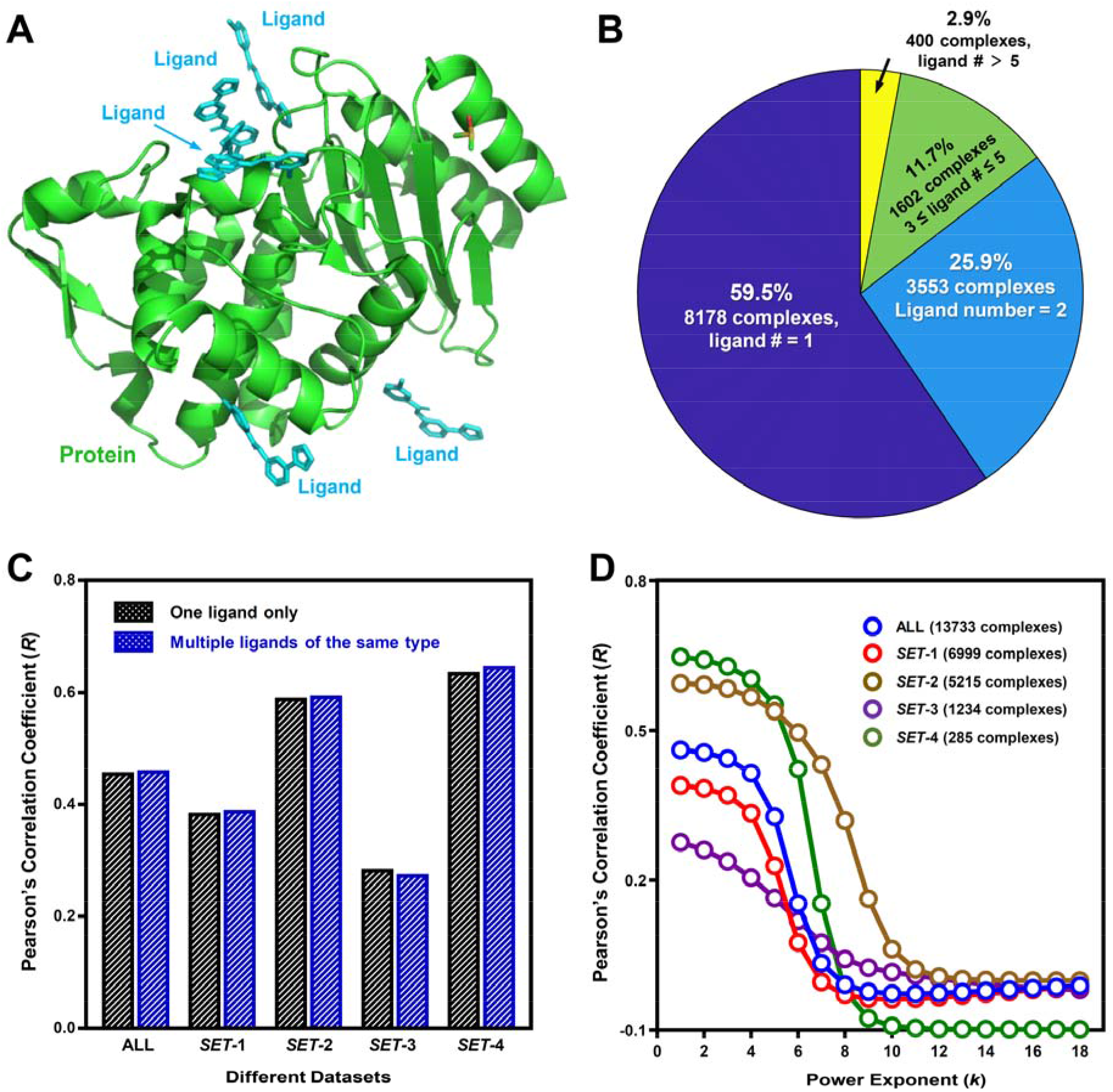
Total number of protein−ligand complexes with multiple binding sites for the same ligand and the performances of untrained models in affinity prediction. A. An example of the experimental structure of the protein−ligand complex with the same ligand bound to multiple sites in the protein. The protein and ligands are colored in green and cyan, respectively. B. Total numbers of protein−ligand complexes with the same ligand bound to one or multiple sites. C. Comparison of the performances of using 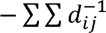 to predict the affinity of one ligand-binding site and using 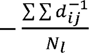 to predict the affinity of the same ligand bound to multiple sites. D. Performances of untrained models, *i.e.*, using the negative sum of the distance-based variables 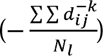 with different power exponents (*k*) to predict the binding affinity.

### Performance of untrained models in binding affinity prediction

The untrained models adopt predefined mathematical formulas for distance-based descriptors without parameter training, where all descriptors are directly extracted from experimental protein–ligand complex structures. The fundamental predefined formulas include the negative summation of distance-based terms: 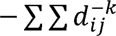 for single-ligand binding systems and 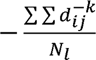 for multi-ligand systems with identical ligands. Here, *d_ij_* represents the distance of the *j*^th^ interacting atom pair belonging to the *i*^th^ atom type pair, *k* denotes the power exponent, and *N_l_* is the total number of bound ligands. These formulas are derived from the classical physical descriptions of electrostatic and van der Waals interaction energies, which are defined as 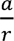 and 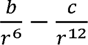 (Levitt, 2001).

When the power exponent (*k*) was set to 1, the Pearson’s correlation coefficient (*R*) values of the single-ligand formula 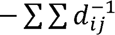 and 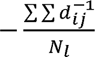 were 0.457 *vs.* 0.461 for the full *ALL* dataset (13,733 complexes), 0.385 *vs.* 0.390 for *SET-1* (6,999 complexes), 0.591 *vs.* 0.594 for *SET-2* (5,215 complexes), 0.285 *vs.* 0.276 for *SET-3* (1,234 complexes), and 0.637 *vs.* 0.647 for *SET-4* (285 complexes) (**Fig. 2C**). Collectively, the multi-ligand calculation strategy yielded slightly improved predictive performance compared with the single-ligand simplification across most datasets, albeit with marginal performance gains.

The predictive performance of the optimized untrained multi-ligand model 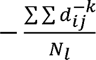 with *k* ranging from 1 to 18 is presented in **Fig. 2D**. For all datasets, *R* values decreased gradually with increasing *k* when *k* ≤ 11, while no substantial performance variation was observed for *k* > 11. Notably, *SET-2* and *SET-4* exhibited superior predictive accuracy relative to *SET-1* and the *ALL* dataset at low power exponents (*k* ≤ 5). This discrepancy can be attributed to data quality differences: *SET-2* and *SET-4* are derived from the high-quality refined subset of PDBbind v.2024, whereas *SET-1* originates from the lower-quality general subset, and the *ALL* dataset integrates data of mixed quality. In contrast, *SET-3* is compiled from the PDBbind v.2024 refined+ subset, which exclusively contains metal-containing protein–ligand complexes. The unique metal-mediated intermolecular interactions in these complexes weaken the correlation between distance-based descriptors and experimental binding affinities, resulting in lower *R* values than other datasets at *k* ≤ 5 (**Fig. 2D**).

### Performance of representative pre-linear and linear models in binding affinity prediction

Based on preliminary predictive performance, 18 representative pre-linear models were screened from a total of 180,000 candidate models. The optimal pre-linear models were obtained at low power exponents (*k* ≤ 3), and a consistent decline in *R* values was observed with increasing *k* for *k* ≥ 4 (**Fig. 3A**). After parameter training, pre-linear models achieved better performance on high-quality datasets (*SET-2, SET-3, SET-4*) than on the low-quality *SET-1* and the mixed-quality *ALL* dataset. Linear regression training significantly improved the predictive power compared with the untrained distance-based summation formulas. Specifically, the *R* values of the optimal untrained model (*k* = 1) and the representative pre-linear model 2 (*k* = 2) were 0.461 *vs.* 0.615 (*ALL*), 0.390 *vs.* 0.558 (*SET-1*), 0.594 *vs.* 0.675 (*SET-2*), 0.276 *vs.* 0.659 (*SET-3*), and 0.647 *vs.* 0.746 (*SET-4*), respectively.

**Figure 3.**
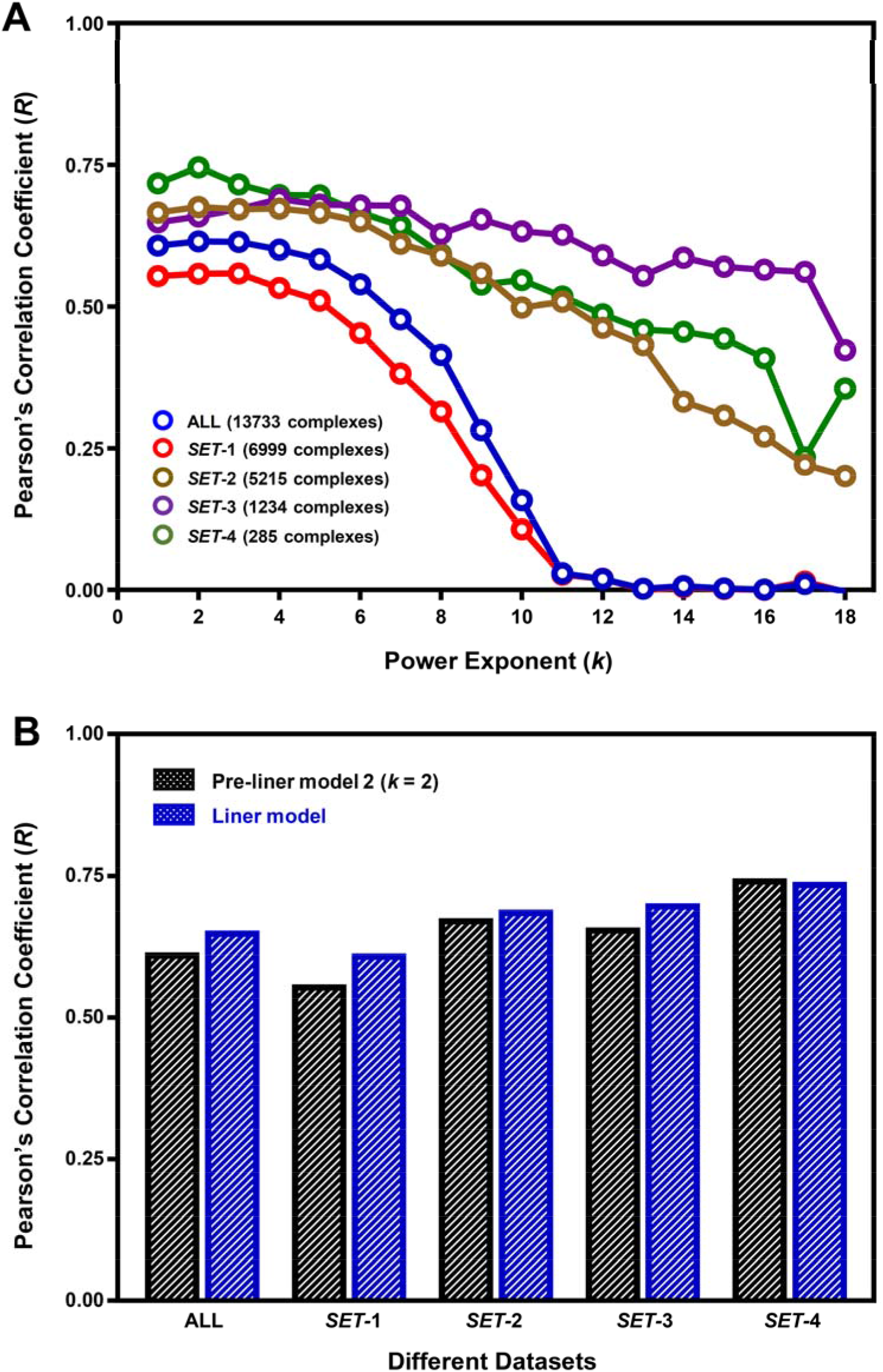
Performances of representative pre-linear models and a representative linear model in affinity prediction. A. Performances of the 18 pre-linear models with different power exponent. B. Comparison of the performances of the pre-linear model 2 (*k* = 2) and the linear model.

To further improve the performances beyond those of the pre-linear models with one single fixed power exponent, a representative linear model was constructed by performing linear regression on the predicted outputs of the 18 pre-screened representative pre-linear models with different power exponents. The performance comparison between pre-linear model 2 (*k* = 2) and the final linear model showed further accuracy improvement: *R* values increased from 0.615 to 0.653 (*ALL*), 0.558 to 0.613 (*SET-1*), 0.675 to 0.690 (*SET-2*), and 0.659 to 0.701 (*SET-3*), with a slight performance drop from 0.746 to 0.739 observed only for *SET-4* (**Fig. 3B**). These results verify that the integration of multi-model predictive outputs effectively enhances the overall predictive performance of linear frameworks.

### Performance of representative neural network and hybrid models in binding affinity prediction

Artificial neural networks were further employed to mine the nonlinear relationship between distance-based descriptors and binding affinity. Eighteen representative artificial neural network models were established for each power exponent, with their average predictive performance summarized in **Fig. 4A**. The artificial neural network models achieved optimal performance at *k* ≤ 5, followed by a steady decrease in *R* values as *k* increased beyond 5. The best-performing artificial neural network model (*k*3-1) yielded *R* values of 0.704 (*ALL*), 0.675 (*SET-1*), 0.732 (*SET-2*), 0.726 (*SET-3*), and 0.800 (*SET-4*), which consistently outperforms all pre-linear and linear models. Note that here, *k* = 3 is the power exponent, and 1 is model number among the 18 representative neural network models using the same original distance-based variables with the same power exponent.

**Figure 4.**
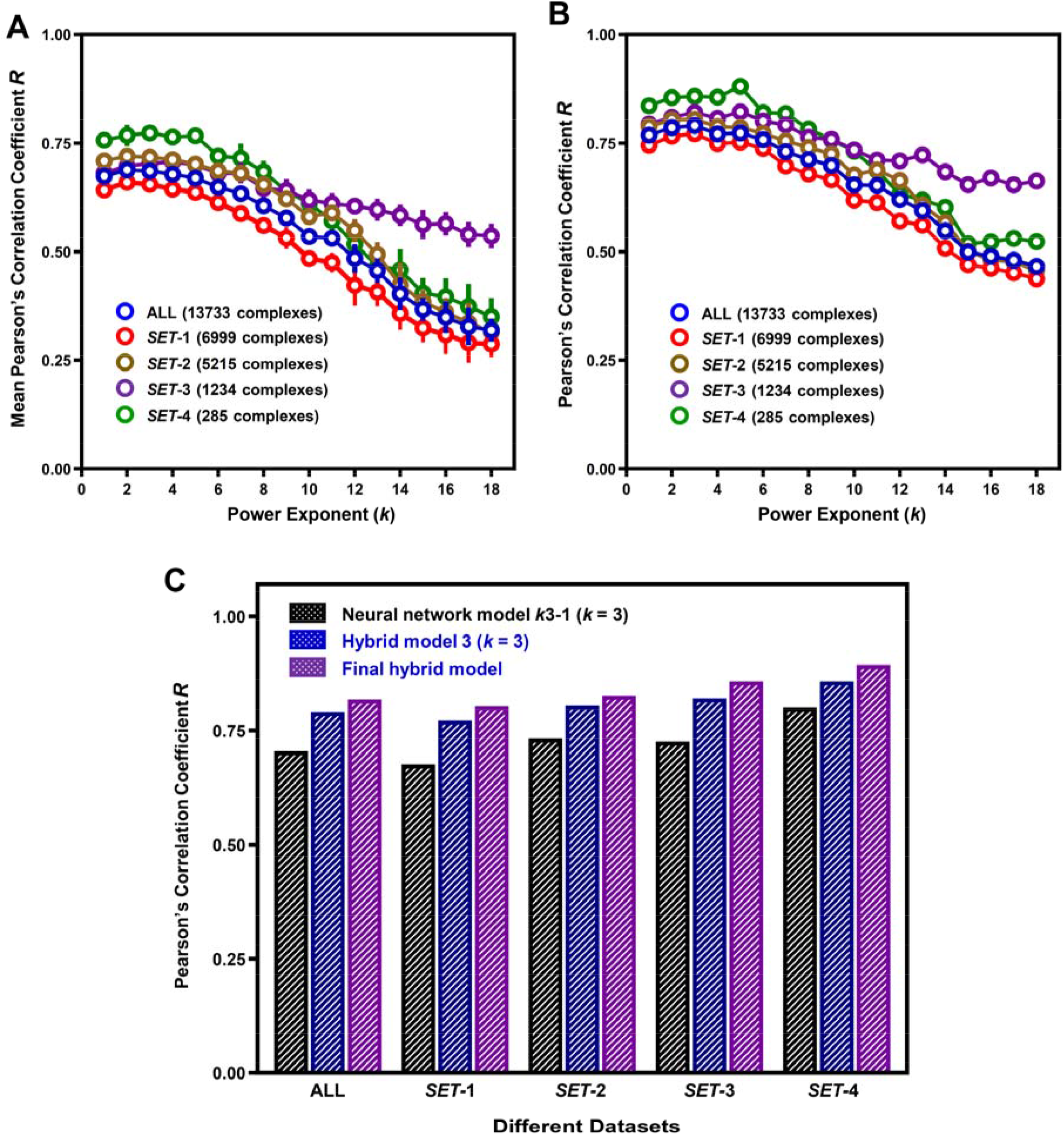
Performances of representative neural network models, representative hybrid models, and final hybrid model in affinity prediction. A. Performances of neural network models with different power exponent. B. Performances of hybrid models with different power exponent. C. Comparison of the performances of neural network model *k*3-1 (*k* = 3), hybrid model 3 (*k* = 3), and final hybrid model.

Subsequently, the hybrid models were developed via linear regression ensemble of artificial neural network model predictions, with one representative hybrid model selected for each power exponent. Across all power exponents, the hybrid models exhibited superior predictive accuracy compared with the corresponding standalone artificial neural network models (**Fig. 4A, 4B**). Consistent with the trends seen with artificial neural network models, hybrid models achieved optimal performance at *k* ≤ 5, with *R* values declining progressively at higher power exponents. Notably, *SET-3* exhibited a much slower performance decay than other datasets. This phenomenon is likely attributed to the unique metal-mediated interactions in *SET-3* complexes, whose interaction strength decays more slowly with increasing power exponents.

Finally, a unified final hybrid model was constructed by linearly integrating the predictive outputs of all 18 representative preliminary hybrid models. The performance progression across the optimal artificial neural network model *k*3-1, preliminary hybrid model (*k* = 3), and final hybrid model is illustrated in **Fig. 4C**. The corresponding *R* values were 0.704, 0.790, and 0.818 (*ALL*); 0.675, 0.772, and 0.803 (*SET-1*); 0.732, 0.804, and 0.826 (*SET-2*); 0.726, 0.821, and 0.858 (*SET-3*); and 0.800, 0.858, and 0.893 (*SET-4*), respectively. The sequential performance improvement across model iterations demonstrates the effectiveness of the hierarchical training and ensemble strategy adopted in this study.

### Comparison with state-of-the-art scoring functions

The scoring and ranking power of the developed linear model, artificial neural network model *k*3-1, hybrid model (*k* = 3), and final hybrid model were benchmarked against conventional and recently reported scoring functions on the CASF-2016 dataset (*SET-4*) (Su et al., 2019). The comparative methods included classical scoring functions (X-Score (Wang et al., 2002), APBScore (Bao et al., 2020), AA-Score (Pan et al., 2022)), a machine learning-based scoring function (Δ_Lin_F9_XGB (Yang and Zhang, 2022)), and eight linear models developed in our previous work (Yang and Zhu, 2025). The performance data of APBScore, AA-Score, and Δ_Lin_F9_XGB were retrieved from their original publications (Bao et al., 2020; Pan et al., 2022; Yang and Zhang, 2022).

As shown in **Fig. 5A, 5B**, all linear scoring functions achieved moderate performance on the CASF-2016 benchmark, with *R* values ranging from 0.63 to 0.76 and mean Spearman’s ρ values ranging from 0.57 to 0.64. In contrast, machine learning-empowered scoring functions (Δ_Lin_F9X_GB, artificial neural network model *k*3-1, hybrid model (*k* = 3), final hybrid model) delivered substantially improved accuracy. Specifically, the Δ_Lin_F9_XGB and final hybrid model achieved *R* values of 0.845 and 0.893, and mean ρ values of 0.704 and 0.800 on CASF-2016, respectively, highlighting the superior predictive capability of the final hybrid model.

**Figure 5.**
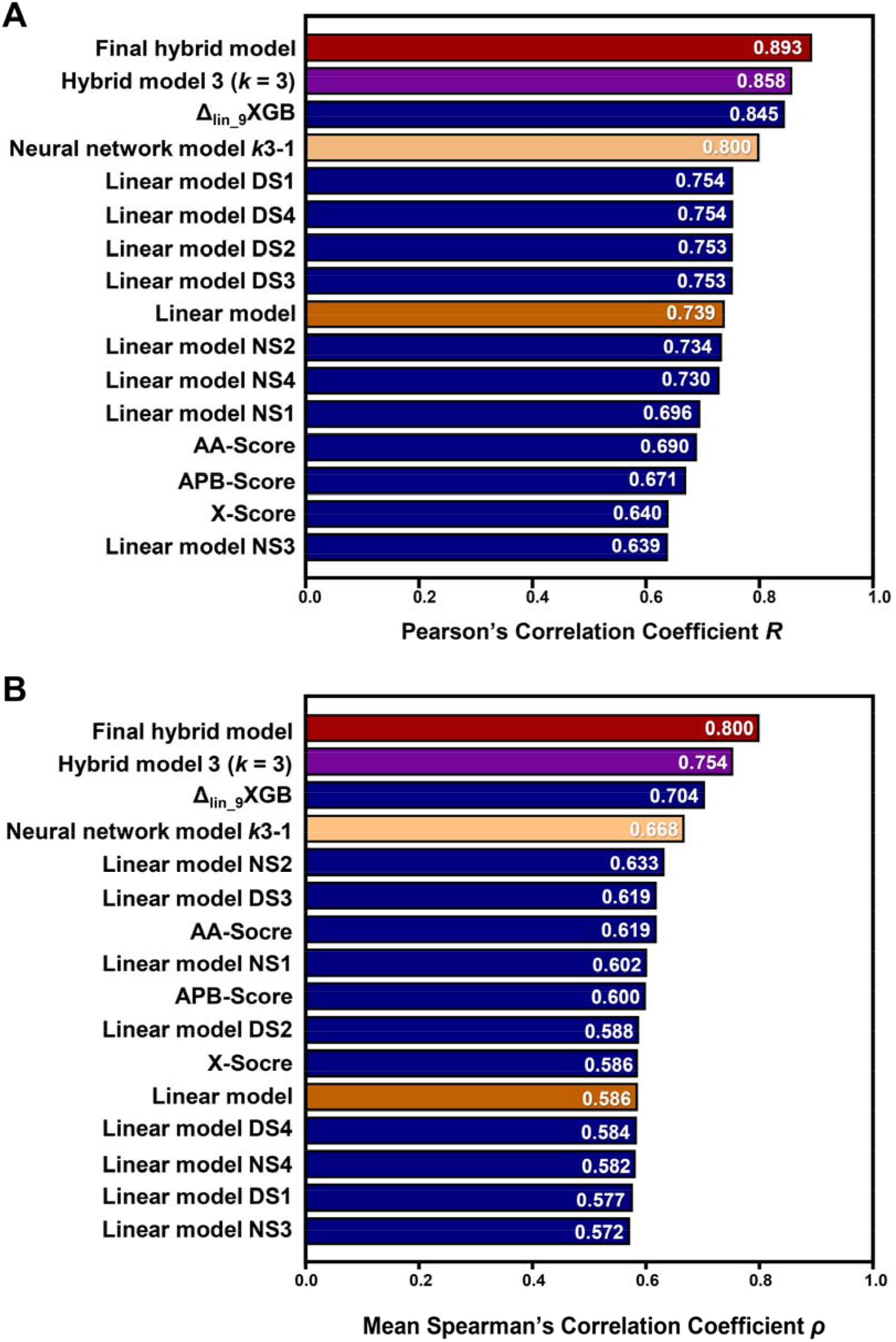
Comparison of four representative models with other state-of-the-art scoring functions in *SET*-4 (CASF-2016). The scoring power (**A**) and ranking power (**B**) of linear model, neural network model *k*3-1 (*k* = 3), hybrid model 3 (*k* = 3), final hybrid model, linear model DS1 DS4, linear model NS1 NS4, X-Score, APBScore, AA-Score, and Δ_Lin_F9_XGB.

### Summary of predictive performance of distance-based models

As quantitatively summarized in **Table 1**, the Pearson’s *R* values across all datasets increased progressively with model complexity, following the order: untrained distance-based model (*k* = 1) → pre-linear model 2 (*k* = 2) → linear model → artificial neural network model *k*3-1 → hybrid model (*k* = 3) → final hybrid model. Similarly, the Spearman’s ρ values on the CASF-2016 (*SET-4*) benchmark also exhibited a consistent upward trend with enhanced model sophistication.

**Table 1.** Performances of the best distance-based models for protein ligand binding affinity prediction.

| Model | Scoring power (Pearson’s correlation coefficient, $R$) | | | | | Ranking power (Spearman’s rank correlation coefficient, $\rho$) |
| --- | --- | --- | --- | --- | --- | --- |
|  | ALL (13733 complexes) | SET-1 (6999 complexes) | SET-2 (5215 complexes) | SET-3 (1234 complexes) | SET-4 (285 complexes) | SET-4 (CASF-2016, 285 complexes) |
| Untrained model (negative sum of the distance-based variables ($k = 1$)) | 0.461 | 0.390 | 0.594 | 0.276 | 0.647 | $0.605 \pm 0.423$ |
| Pre-linear model 2 ($k = 2$) | 0.615 | 0.558 | 0.675 | 0.659 | 0.746 | $0.577 \pm 0.363$ |
| Linear model | 0.653 | 0.613 | 0.690 | 0.701 | 0.739 | $0.586 \pm 0.359$ |
| Neural network model $k3-1$ | 0.704 | 0.675 | 0.732 | 0.726 | 0.800 | $0.668 \pm 0.326$ |
| Hybrid model ($k = 3$) | 0.790 | 0.772 | 0.804 | 0.821 | 0.858 | $0.754 \pm 0.256$ |
| Final hybrid model | 0.818 | 0.803 | 0.826 | 0.858 | 0.893 | $0.800 \pm 0.229$ |

Consistent with our previous findings (Yang and Zhu, 2025), the present results demonstrate that linear models trained on high-quality affinity datasets can achieve *R* values exceeding 0.700. To further validate this conclusion, a high-quality dataset consisting of 5,500 complexes from *SET-2* and *SET-4* (covering 651 atom type pairs) was constructed, with 3,500 complexes randomly selected as the training set for supplementary pre-linear and linear model development. The optimized supplementary linear model achieved *R* values of 0.731 (*SET-2*) and 0.802 (*SET-4*), with a mean ρ value of 0.677 ± 0.327 on CASF-2016 (**Supplementary Table S1**). This performance is comparable to or even superior to several advanced machine learning-based scoring functions, including Δ_vina_RF_20_ (*R* = 0.732, ρ = 0.626) (Wang and Zhang, 2017) and Δ_vina_XGB (*R* = 0.796, ρ = 0.647) (Lu et al., 2019) on the CASF-2016 benchmark.

Collectively, the optimized linear models based on distance-based descriptors achieved *R* values of 0.700–0.800 on high-quality datasets, while the hybrid artificial neural network-based models further improved the accuracy to 0.800–0.900.

## DISCUSSION

### Data leakage and over fitting

The data leakage and over fitting are the common problems in the machine learning-based and deep learning-based models, but they are not completely avoided in practice. The data similarity between the different sets is the key for a model to work in other sets beyond the training data. In terms of protein ligand binding affinity prediction, it is no other than the interaction similarities among the structures in different sets which make the strong generalization ability of the scoring functions/models possible. Indeed, the lack of completely independent test sets is definitely the key defection of this work. In order to the explore the upper boundary of predictive powers of distance-based models in protein ligand binding affinity prediction, the models are trained, integrated and selected by fair means or foul. This work has not yet provided the conclusive models. With the increasing of high-quality data, the models will be improved and the conclusive models will be selected and tested in different external datasets.

The computational prediction accuracy for protein ligand binding affinity is, in theory, lower than the precision based on experimental measurements (*R* ≈ 0.900) (Kramer et al., 2012). The overfitting criteria should be the ultrahigh performances (*R* > 0.900, or even ≈ 1) in different sets. In this work, the *R* value of the untrained model in *SET*-4 (CASF-2016) can achieve to 0.647 (**Table 1**), the *R* value of the best linear model achieves to 0.802 (**Supplementary Table S1**), and the *R* value of the final complicated model achieves to 0.893 (**Table 1**), which does not exceed the expectation. The representative models were shown in this work to exhibit the possible upper boundary of the best performances of distance-based protein ligand binding affinity prediction.

While the training-overlap result analysis in this exploration did not strictly adopt the common practices in data science, the findings are still meaningful and inspiring for future research endeavors in this or related areas. It is expected that the performances of linear models with considerable generalization ability will further improve if the improved dataset can include over 2200 complexes with 108 kinds of non-sparse interacting atom-type pairs and each kind of atom-type pair exists in over 300 protein–ligand complexes. As the *R* values between the predicted and experimental binding affinities in subsets composed of selected complexes are between 0.850 and 0.900 (**Supplementary Table S2**), it is hopeful to uncover the precise scoring function with an explicit mathematical expression that enables the calculation of binding affinity with even higher accuracy. The affinity prediction within the predicted experimental error (*R* ≈ 0.900) (Kramer et al., 2012) is still of great practical value for protein–ligand interactions only involving the known kinds of atom-type pairs in the scoring functions.

In contrast to the abundant sequence and structural data available to support AlphaFold2 (Jumper et al., 2021) development, high-quality experimental affinity data paired with high-resolution protein–ligand complex structures remain scarce, which restricts the training of high-performance deep learning models for affinity prediction. It is expected that the predictive *R* values may further improve with increasing model complexity and decreasing model generalization ability, however, such improvements may not be as significant or useful in practical applications with unknown or new datasets. The balance between the complexity and the performance should follow the rule of Ockham’s razor. Theoretically, a perfect scoring function or model should not be influenced by the training and test sets, and should exhibit high prediction accuracies in any datasets. Although the other methods compared with the models in this work were not trained based on the newest data, without regard to the problems of data leakage and over fitting, at least they should be successful in the previous existed sets such as CASF-2016 Core Set (*SET*-4) (Su et al., 2019). At all events, the perfect scoring function remains to be discovered in the future.

### Model interpretability and physical significance

Generally, the interpretability of the models with explicit mathematical expressions such as the traditional empirical scoring functions (Bao et al., 2020; Pan et al., 2022; Wang et al., 2002; Xie et al., 2025; Yang and Zhang, 2021) is better than that of the models without explicit mathematical expressions (Lu et al., 2019; Su et al., 2019; Wang and Zhang, 2017; Wang et al., 2002; Yang and Zhang, 2022). However, the performances of the traditional empirical scoring functions (0.600 < *R* < 0.700) are usually worse than those of the machine learning-based scoring functions (0.700 < *R* < 0.900). Similarly, the interpretability of the pre-linear and linear models in this work (0.600 < *R* < 0.800) are better than that of the neural network, hybrid and final hybrid models (0.700 < *R* < 0.900).

After breaking the restrictions in the traditional atomic interaction energy terms, the affinity predictive power of the scoring functions with explicit mathematical expressions was improved by determining the atom type pairs’ coefficients simultaneously. In the background of protein ligand interactions, the effect of multiple atomic interactions may be implicitly integrated into the scoring functions, which can’t be fully reflected in the traditional empirical functions based on the individual and intact energy and geometric terms. The optimal performances with lower power exponents (k ≤ 5) indicate that the decay shape of the distance-based interactions is not extremely steep and precipitous. Although taking no account of the contributions of protein ligand binding process to binding affinity apparently, the scoring functions with different power exponents should implicitly incorporate the most factors about binding such as solvent effects and conformational entropy. Different atom-type pairs with different properties make different contributions to the binding affinity, which would be reflected on the different coefficients in the scoring functions. Although there are no explicit mathematical expressions for the neural network, hybrid and final hybrid models, the effect of each interacting atom pair can still be estimated by altering the original distance from the original value to a smaller or bigger one. With sufficient training and test data, the generated models could also reflect the effect of mutation accurately. The physical significance of the models with expanded power exponents will be deeply explored and completed in the future.

The protein ligand binding affinity concerns the binding strength which is the key of scoring and ranking powers. The screening power refers to the problem about binding or not, and the docking power is a structural prediction problem. Although an ideal scoring function should be versatile in different powers, the scoring functions in practice often failed to be professional in different problems simultaneously. In the future, the conclusive models for affinity prediction will be applied in the docking and screening fields.

Additionally, the linear and final hybrid models with one single fixed power exponent are superior to the pre-linear, neural network and hybrid models with multiple power exponents (**Table 1** and **Supplementary Table S1**). The comparable performances of the models shown in this work can be reproduced in the given sets based on the description in **Materials and Methods** section. But there is no need to generate the completely same models using the same machine learning method and training parameters. It is very possible that the other frequently-used machine learning methods such as random forest can also make the comparable performances.

### Future outlook

Although the mathematical formulations of electrostatic and van der Waals interaction energies have been adopted as essential terms in scoring functions for affinity prediction and can serve as valid theoretical references, it remains challenging to derive an optimal mathematical formulation with superior predictive performance based solely on a single structural snapshot of protein–ligand complexes. This study only benchmarks the upper predictive limit of distance-based descriptors via artificial neural network and linear regression methods, without developing an explicit, optimally performing mathematical expression for affinity prediction. Although artificial neural network models lack explicit analytical expressions, their outstanding predictive performance implies the existence of potential universal mathematical formulas capable of precise protein–ligand binding affinity prediction. Future research can build on the growing volume of high-quality experimental data to further explore, excavate and validate the intrinsic distance–energy correlation with explicit mathematical expressions governing protein–ligand interactions, which will be crucial for advancing the accuracy and interpretability of affinity prediction models. Furthermore, the developed distance-based scoring functions will be applied to binding pose selection and large-scale structural prediction of protein–ligand complexes using experimental structural datasets.

According to the research and practice in this work, the problems to be handled in the future are summarized as follows:

### □. Accurately determination of the atom type pairs’ coefficients

The atom type pairs’ coefficients may be not very accurate because of the inaccuracy of sparse matrix’s inverse solved by numerical calculation. For example, the number of variables in the supplementary pre-linear models are same, *i.e.*, 651, and the dimension of the matrix is 652 (651 kinds of atom type pairs + 1 constant term). But the performances of supplementary pre-linear model 11 (power exponent *k* = 11) (**Supplementary Table S1**) are much better than those of supplementary pre-linear model 12 (power exponent *k* = 12). The reason may be that the matrix for determining the coefficients in supplementary pre-linear model 12 is sparser with the much smaller values of the distance-based variables than that for the supplementary pre-linear model 11 (power exponent *k* = 11). This makes the more inaccuracy of the atom type pairs’ coefficients.

Additionally, only one single fixed exponent was adopted in the pre-linear models, which restricts the improvement of affinity prediction accuracy. In the other supplementary linear models generated based on the data without sparse atom type pairs (**Supplementary Table S2**), the number of power exponents were not restricted, but the performances of supplementary linear model 12 (12 adopted power exponents, dimension *n* = 1297 = 12×108 variables + 1 constant term) are much worse than those of supplementary linear model 11 (11 adopted power exponents, dimension *n* = 1189 = 11×108 variables + 1 constant term). One possible reason is the inaccuracy of high-dimensional (dimension *n* > 1200) or sparse matrix’s inverse solved by numerical calculation when determining the atom type pairs’ coefficients.

Another noteworthy issue is that the performances (0.850 < *R* < 0.900) of the supplementary linear model 10 (**Supplementary Table S2**) can’t be maintained by the newly-added or determined coefficients for the emerging interacting atom-type pairs in the expanded datasets. The newly-added pairs exist in fewer than 300 protein–ligand complexes, which may lead to non-optimal coefficients induced by inaccuracy of the inverse of the sparse matrix. Another reason is that the optimized groups of coefficients in previous linear models for the emerging pairs are not picked from different groups of coefficients with the best performances. In the future, with increasing high-quality data, it is worthwhile to further explore the scoring functions with explicit mathematical expressions to achieve and maintain best performances in the expanded datasets.

Another alternative way to determine the coefficients is using optimization algorithm rather than least-squares method (Nievergelt, 2000). The sparse atom type pairs could be clustered and merged into several classes. Certainly, the traditional electrostatic and van der Waals energy terms can be directly employed to characterize the interactions of sparse atom type pairs. These ways are also worth to be explored in the future.

### □. Implicit consideration of the metal ions, water molecules, multibody atomic interactions

Indeed, the metal ions should be considered specifically in the metal-involving protein ligand interactions. However, only 1234 metal-containing protein ligand systems exist in *SET*-3. The receptor metal ion ligand atom type pairs would be very sparse in the set. In this work, the metal ions-mediated, water molecule-mediated and the multibody atomic interactions between protein and ligand were only implicitly considered in the scoring functions and models. In the next step, we will consider the metal ions explicitly and detailly.

### □. Adequate consideration of the ligand number and size

Multiple ligands with same type exist in the structures of protein ligand complexes, even in the same specific binding pocket. In the beginning, this work intended to improve the affinity predictive power of distance-based variables by considering the multiple ligands with same type in the structures of protein ligand complexes. We employed the sum of the interactions of the same ligand at different binding sites and the corresponding magnification based on the experimental binding affinity, and the correlations between them would increase dramatically when comparing with those in the only-one-ligand-considered situation. However, this may be caused by the different numbers of ligands in different complexes which stretches the data points into a long and narrow region. The averaging approach is the simple one to consider the multiple ligands with same type in the complexes. But the improvement is not apparent. In the future, we will consider more reasonable ways to handle this problem.

When making interactions with the protein receptors, the small ligands may be different from the large ones such as peptides and protein monomers. Although a scoring function should adapt to all the protein ligand systems theoretically, in order to decrease the influence of ligand size, only small ligands were considered in this work. It is expected that the performance will decrease when predicting the large ligand protein binding affinities. The effect of ligand size will be considered and investigated in the future.

### □. Construction of external independent test sets

Except for the PDBbind dataset which collected the public experimental structural and affinity data (Liu et al., 2017; Su et al., 2019; Wang et al., 2004), no enough external experimental structures with reliable binding affinity constitute the new test set which should be composed of at least several hundred high-quality data. Although there exist a large number of modelled structures with experimental binding affinity in BindingNet (Li et al., 2024; Zhu et al., 2025), these modelled structures may have deviations from the experimental ones. Additionally, the selecting process of the representative models in this study was not very rigorous. This work explores the upper boundary of the predictive power of distance-based protein ligand binding affinity prediction, but the training-overlap test makes the results not extremely solid. In the future, the models will be modified and improved, and new high-quality data would be adopted as the external test set, and the modelled structures would also be considered in the training and independent test sets.

## CONCLUSION

In traditional force fields, nonbonded atomic interactions, including electrostatic interactions, van der Waals forces, and hydrogen bonding, are predominantly distance-dependent. In this study, we further explored and optimized the predictive performance of distance-based variables for protein–ligand binding affinity prediction via linear regression and artificial neural network algorithms. By extending the power exponents of distance terms, the established scoring functions implicitly integrate intrinsic multibody interaction effects. In some instances, the optimal models developed in this work outperform previously reported state-of-the-art prediction methods. Specifically, the results give the hope that the scoring functions with explicit mathematical expressions can achieve a near-experimental-error affinity prediction accuracy after appropriately handling the problems of the sparse atom-type pair coefficients and accurately solving the high-dimensional matrix inverse (dimension *n* > 1200) by numerical calculation. Overall, the findings validate the efficacy and potential of refining distance-based variable descriptors to enhance the accuracy of protein–ligand binding affinity prediction.

## MATERIALS AND METHODS

### Datasets

The protein–ligand binding affinity data used in this study were curated from PDBbind (version 2024) (Liu et al., 2017; Wang et al., 2004). The corresponding three-dimensional complex structures were retrieved from the Protein Data Bank (https://www.rcsb.org/) (Berman et al., 2000). All complexes were filtered according to the following five criteria: ***i.*** Valid experimental binding affinity values (*i.e.*, inhibition constant *K* or dissociation constant *K*_d_) were available. ***ii.*** Each ligand contained no more than 150 total atoms and 70 heavy atoms. ***iii.*** No steric clashes between heavy atoms, defined as an interatomic distance greater than 1 Å between any two heavy atoms. ***iv.*** Protein–peptide complexes were excluded. ***v.*** Binding pockets contained no atypical or unrecognized atom types, including Cd, Cu, As, Y, Yb, Ru, Pt, V, Si, Be, Al, Rb, Pr, Mo, Cs, Te, Sr, Re, In, Ga, Eu and Du. Each of the atypical atom types exists in no more than 5 complexes. After filtering, a total of 13,733 protein–ligand complexes were retained. These samples were further partitioned into four subsets following the data quality classification of PDBbind v2024 (Liu et al., 2017; Wang et al., 2004): *SET-1* (6,999 complexes) from the PDBbind general set; *SET-2* (5,215 complexes) from the high-quality PDBbind refined set; *SET-3* (1,234 complexes) from the refined+ set consisting of high-quality complexes and metal-binding protein–ligand systems; and *SET-4* (285 complexes), corresponding to the CASF-2016 Core Set (Su et al., 2019).

### Atom types and interacting pairs

Atom typing for all experimental structures was performed using Open Babel 2.4.1 (O’Boyle et al., 2011). In total, 36 types of heavy atoms were identified within the binding pockets of both proteins and ligands (**Supplementary Table S3**). Water molecules were removed prior to subsequent analysis, while multiple identical ligands in a single structure were preserved. Consistent with a previous study (Yang and Zhu, 2025), an interacting atom pair was defined as two atoms originating separately from the ligand and the protein receptor, with an interatomic distance ranging from 1 Å to 10 Å. Notably, atom pairs were directionally distinguished: a pair consisting of ligand atom type 1 and receptor atom type 2 was treated as distinct from the reverse combination (ligand atom type 2 and receptor atom type 1), provided the two atom types differed. Ultimately, 859 unique directional atom-type pairs were obtained (**Supplementary File 1**).

### Affinity prediction models

#### Pre-linear and linear models

Distance-based pre-linear models were constructed via the least-squares method (Nievergelt, 2000), with the mathematical formulation presented below:

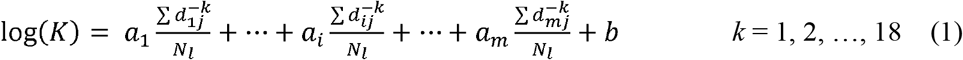

where *K* denotes either the inhibition constant (*K*_i_) or dissociation constant (*K*_d_); the experimental *K*□ or *K*_d_; *a*□ and *b* represent regression coefficients and the intercept term, respectively; *N_l_* is the number of identical ligands in one complex; *d_ij_* refers to the distance of the *j*^th^ interacting pair belonging to the *i*^th^ atom-type pair; *m* equals 859 (the total number of atom-type pairs); and *k* is the power exponent, where each value of *k* corresponds to an independent pre-linear model.

A training set of 8,000 complexes was randomly sampled from the full dataset (13,733 complexes), covering all 859 atom-type pairs. This random sampling was repeated 10,000 times to generate 10,000 independent training sets for model training. One representative pre-linear model was selected for each power exponent from the resulting 180,000 preliminary models.

Linear models were further built based on the optimized pre-linear models, as described in Equation (2):

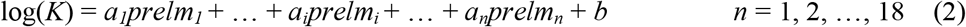

where *K*, *a*□ and *b* follow the same definitions as above; *prelm_i_* is the predicted value from the *i*^th^ representative pre-linear model; *n* indicates the maximum power exponent of the adopted distance-based variables.

Training sets for linear models were generated using the same random sampling strategy as for pre-linear models, with 100 replicates. All possible combinations of the 18 pre-linear models (2^18^ - 1) combinations) were adopted for linear model construction, yielding a total of 26,214,300 linear models ((2^18^ - 1) × 100). A single optimal linear model was chosen according to its predictive performance across the four datasets.

#### Neural network and hybrid models

To further explore the predictive capability of distance-based descriptors, backpropagation neural networks (BPNN) (Li et al., 2012; Rumelhart et al., 1986) were developed using the same input variables as the pre-linear models. The network architectures contained either one or two hidden layers, with node numbers per layer set to 5, 10, 15 or 20. This design produced 20 distinct network configurations (4 + 4 × 4). For each architecture, 10 independent models were trained to optimize network parameters, resulting in 200 models in total. Eighteen representative neural network models (one per power exponent) were selected, using the training set applied for pre-linear model optimization.

First-stage hybrid models were constructed by integrating representative neural network models with the same power exponent, following Equation (3):

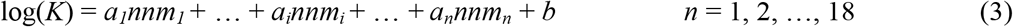

where *nnm_i_* is the predicted value using the *i^th^* neural network model, and other parameters are defined as previously described. The training sets were consistent with those used for linear hybrid models derived from pre-linear models. One representative hybrid model was selected for each power exponent based on performance across the four datasets.

Finally, second-stage integrated hybrid models were established using the 18 first-stage hybrid models, with the formula shown in Equation (4):

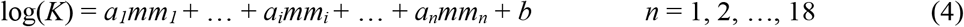

where *mm_i_* represents the predicted value from the *i*^th^ first-stage hybrid model. The same training sets were utilized here. The ultimate representative hybrid model was determined by its performance on all four subsets.

### Model evaluation metrics

The scoring power of prediction models was evaluated using the Pearson’s correlation coefficient (*R*) (Su et al., 2019), calculated as:

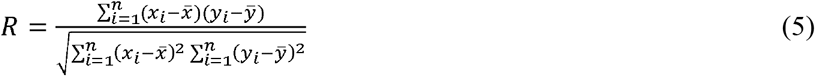

where *n* is the total number of complexes in a dataset; *x_i_* and *y_i_*denote experimental and predicted binding affinities of the *i*^th^ complex, respectively; *x̅* and *y̅* are the corresponding mean values.

Model ranking ability was assessed via the Spearman’s rank correlation coefficient (ρ) (Spearman, 1904; Su et al., 2019):

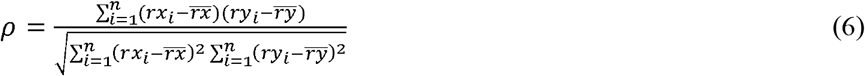

where *n* refers to the number of complexes for a single protein target and binding pocket; *rx_i_* and *ry_i_* are the ranks of experimental and predicted affinities for the *i*^th^ complex; *r̅x̅* and *r̅y̅* are the average ranks. The CASF-2016 Core Set (*SET-4*) contains 57 unique protein targets, each bound by 5 distinct ligands (Su et al., 2019). The overall ranking performance was quantified by the average ρ value across all 57 targets.

## Data availability statement

**Supplementary File 1.** The 859 kinds of atom type pairs and the frequencies of these atom type pairs among the 13733 complexes. **Supplementary File 2.** The negative sums of the distance-based variables. **Supplementary File 3.** The coefficients of the representative pre-linear models, linear model, hybrid models and final hybrid model. **Supplementary File 4.** The predicted values of all the representative models in the results section. **Supplementary File**The performances of all the representative models in the results section. **Supplementary File** The 651 kinds of atom type pairs and the frequencies of these atom type pairs among the 5500 complexes. **Supplementary File 7.** The 108 kinds of atom type pairs and the frequencies of these atom type pairs among the 2227 complexes. **Supplementary Table S1.** Performances of the supplementary pre-linear and linear models using the high-quality affinity data. **Supplementary Table S2.** Performances of the supplementary linear models using the structural data without sparse atom type pairs. **Supplementary Table S3.** Definition of the heavy atom types and the frequencies of these atom types among the 13733 complexes.

The supplemental materials could be downloaded from <u>10.5281/zenodo.23434644</u>. All the data of the study are available upon request.

## CONFLICT OF INTEREST

The authors declare that they have no conflict of interest.

## AUTHOR CONTRIBUTIONS

Conceptualization, Y.X.Y.; Research design: Y.X.Y.; Methodology, Y.X.Y.; Formal analysis, Y.X.Y., B.T.Z.; Investigation, Y.X.Y., B.T.Z.; Resources, B.T.Z.; Data curation, Y.X.Y.; Writing—original draft preparation, Y.X.Y., B.T.Z.; Writing—review and editing, Y.X.Y., B.T.Z.; Visualization, Y.X.Y.; Supervision, B.T.Z.; Project administration, B.T.Z.; Funding acquisition, B.T.Z.

## ACKNOWLEDGEMENTS

This work is supported by research grants from Shenzhen Key Laboratory of Steroid Drug Discovery and Development (No. ZDSYS20190902093417963) and Shenzhen Peacock Plan (No. KQTD2016053117035204).

---

**Supplementary Table S1.**
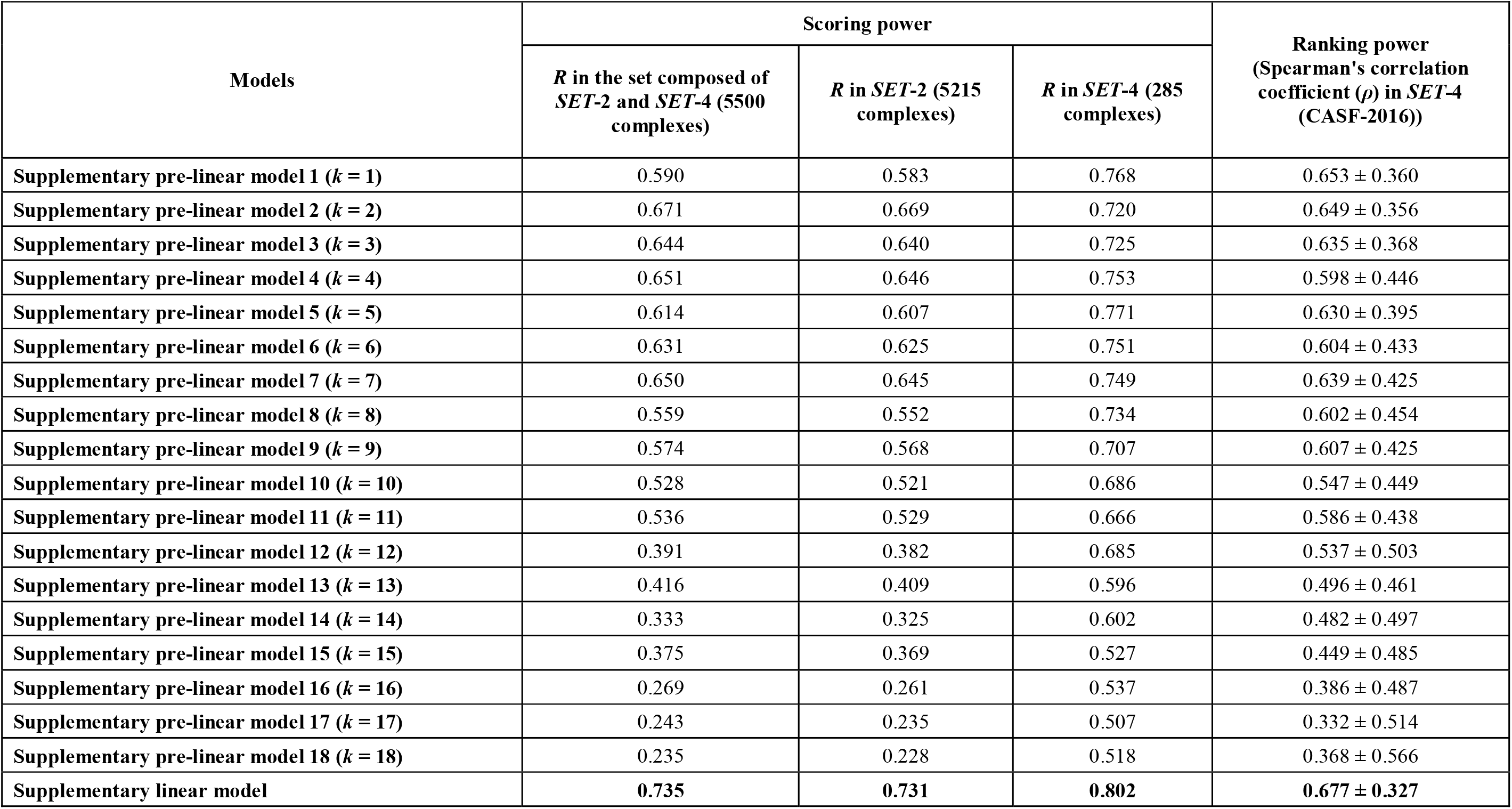
Performances of the supplementary pre-linear and linear models using the high-quality affinity data.

| Models | Scoring power | | | Ranking power<br>(Spearman's correlation coefficient ($\rho$) in *SET-4* (CASF-2016)) |
| --- | --- | --- | --- | --- |
|  | *R* in the set composed of *SET-2* and *SET-4* (5500 complexes) | *R* in *SET-2* (5215 complexes) | *R* in *SET-4* (285 complexes) |  |
| Supplementary pre-linear model 1 ($k = 1$) | 0.590 | 0.583 | 0.768 | $0.653 \pm 0.360$ |
| Supplementary pre-linear model 2 ($k = 2$) | 0.671 | 0.669 | 0.720 | $0.649 \pm 0.356$ |
| Supplementary pre-linear model 3 ($k = 3$) | 0.644 | 0.640 | 0.725 | $0.635 \pm 0.368$ |
| Supplementary pre-linear model 4 ($k = 4$) | 0.651 | 0.646 | 0.753 | $0.598 \pm 0.446$ |
| Supplementary pre-linear model 5 ($k = 5$) | 0.614 | 0.607 | 0.771 | $0.630 \pm 0.395$ |
| Supplementary pre-linear model 6 ($k = 6$) | 0.631 | 0.625 | 0.751 | $0.604 \pm 0.433$ |
| Supplementary pre-linear model 7 ($k = 7$) | 0.650 | 0.645 | 0.749 | $0.639 \pm 0.425$ |
| Supplementary pre-linear model 8 ($k = 8$) | 0.559 | 0.552 | 0.734 | $0.602 \pm 0.454$ |
| Supplementary pre-linear model 9 ($k = 9$) | 0.574 | 0.568 | 0.707 | $0.607 \pm 0.425$ |
| Supplementary pre-linear model 10 ($k = 10$) | 0.528 | 0.521 | 0.686 | $0.547 \pm 0.449$ |
| Supplementary pre-linear model 11 ($k = 11$) | 0.536 | 0.529 | 0.666 | $0.586 \pm 0.438$ |
| Supplementary pre-linear model 12 ($k = 12$) | 0.391 | 0.382 | 0.685 | $0.537 \pm 0.503$ |
| Supplementary pre-linear model 13 ($k = 13$) | 0.416 | 0.409 | 0.596 | $0.496 \pm 0.461$ |
| Supplementary pre-linear model 14 ($k = 14$) | 0.333 | 0.325 | 0.602 | $0.482 \pm 0.497$ |
| Supplementary pre-linear model 15 ($k = 15$) | 0.375 | 0.369 | 0.527 | $0.449 \pm 0.485$ |
| Supplementary pre-linear model 16 ($k = 16$) | 0.269 | 0.261 | 0.537 | $0.386 \pm 0.487$ |
| Supplementary pre-linear model 17 ($k = 17$) | 0.243 | 0.235 | 0.507 | $0.332 \pm 0.514$ |
| Supplementary pre-linear model 18 ($k = 18$) | 0.235 | 0.228 | 0.518 | $0.368 \pm 0.566$ |
| Supplementary linear model | **0.735** | **0.731** | **0.802** | **$0.677 \pm 0.327$** |

**Supplementary Table S2.**
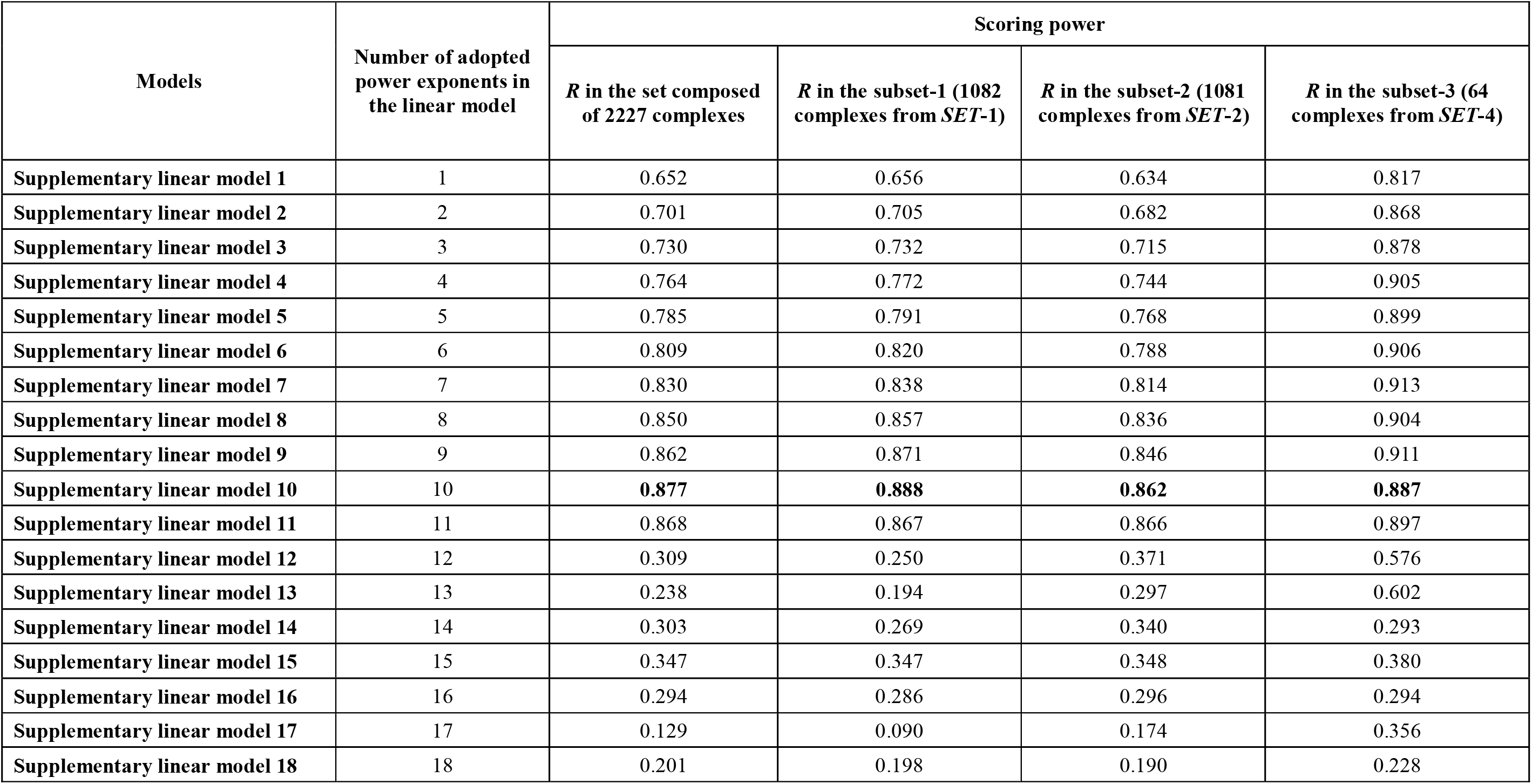
Performances of the supplementary linear models using the structural data without sparse atom type pairs.

**Supplementary Table S3.**
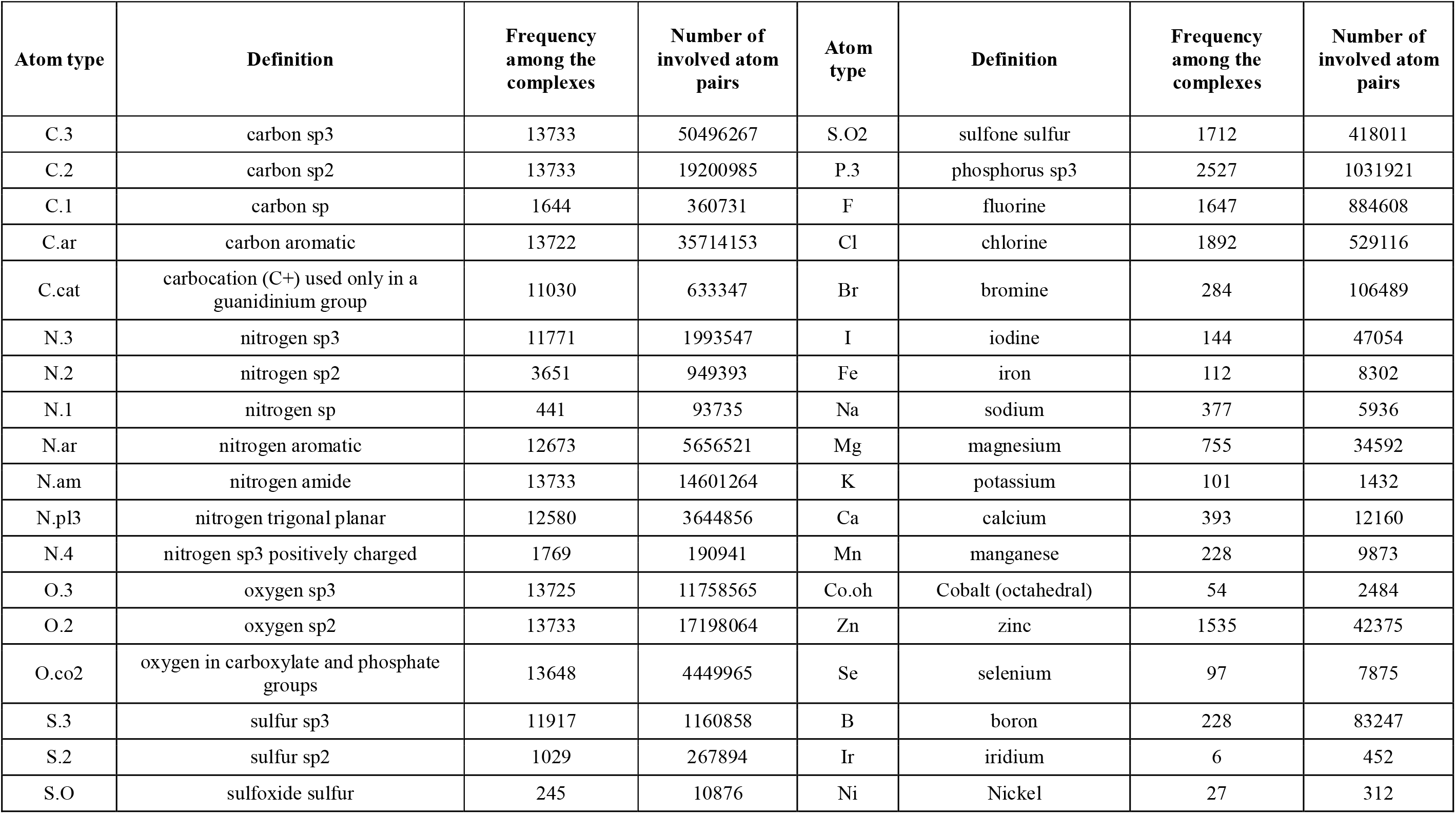
Definition of the heavy atom types and the frequencies of these atom types among the 13733 complexes.

## Notes

1 **Funding Support:** This work was supported, in part, by research grants from Shenzhen Peacock Plan (No. KQTD2016053117035204) and Shenzhen Key Laboratory of Steroid Drug Discovery and Development Research (No. ZDSYS20190902093417963).

### Competing Interest Statement

The authors have declared no competing interest.

### Summary of Updates

More detail were wrote in the revised manuscript and the supplementary files and Tables.

