## Supplementary material for "Exploring the Upper Boundaries of Distance-Based Protein–Ligand Binding Affinity Prediction: Accuracy Improvements via Linear Regression and Artificial Neural Networks": Supplementary Table S1 Performances of the supplementary pre-linear and linear models using the high-quality affinity data.docx

| **Models** | **Scoring power** | | | **Ranking power**  **(Spearman correlation coefficient (*ρ*) in *SET*-4 (CASF-2016))** |
| --- | --- | --- | --- | --- |
|  | ***R* in the set composed of *SET*-2 and *SET*-4 (5500 complexes)** | ***R* in *SET*-2 (5215 complexes)** | ***R* in *SET*-4 (285 complexes)** |  |
| **Supplementary pre-linear model 1 (*k* = 1)** | 0.590 | 0.583 | 0.768 | 0.653±0.360 |
| **Supplementary pre-linear model 2 (*k* = 2)** | 0.671 | 0.669 | 0.720 | 0.649±0.356 |
| **Supplementary pre-linear model 3 (*k* = 3)** | 0.644 | 0.640 | 0.725 | 0.635±0.368 |
| **Supplementary pre-linear model 4 (*k* = 4)** | 0.651 | 0.646 | 0.753 | 0.598±0.446 |
| **Supplementary pre-linear model 5 (*k* = 5)** | 0.614 | 0.607 | 0.771 | 0.630±0.395 |
| **Supplementary pre-linear model 6 (*k* = 6)** | 0.631 | 0.625 | 0.751 | 0.604±0.433 |
| **Supplementary pre-linear model 7 (*k* = 7)** | 0.650 | 0.645 | 0.749 | 0.639±0.425 |
| **Supplementary pre-linear model 8 (*k* = 8)** | 0.559 | 0.552 | 0.734 | 0.602±0.454 |
| **Supplementary pre-linear model 9 (*k* = 9)** | 0.574 | 0.568 | 0.707 | 0.607±0.425 |
| **Supplementary pre-linear model 10 (*k* = 10)** | 0.528 | 0.521 | 0.686 | 0.547±0.449 |
| **Supplementary pre-linear model 11 (*k* = 11)** | 0.536 | 0.529 | 0.666 | 0.586±0.438 |
| **Supplementary pre-linear model 12 (*k* = 12)** | 0.391 | 0.382 | 0.685 | 0.537±0.503 |
| **Supplementary pre-linear model 13 (*k* = 13)** | 0.416 | 0.409 | 0.596 | 0.496±0.461 |
| **Supplementary pre-linear model 14 (*k* = 14)** | 0.333 | 0.325 | 0.602 | 0.482±0.497 |
| **Supplementary pre-linear model 15 (*k* = 15)** | 0.375 | 0.369 | 0.527 | 0.449±0.485 |
| **Supplementary pre-linear model 16 (*k* = 16)** | 0.269 | 0.261 | 0.537 | 0.386±0.487 |
| **Supplementary pre-linear model 17 (*k* = 17)** | 0.243 | 0.235 | 0.507 | 0.332±0.514 |
| **Supplementary pre-linear model 18 (*k* = 18)** | 0.235 | 0.228 | 0.518 | 0.368±0.566 |
| **Supplementary linear model** | **0.735** | **0.731** | **0.802** | **0.677±0.327** |
