## Supplementary material for "Exploring the Upper Boundaries of Distance-Based Protein–Ligand Binding Affinity Prediction: Accuracy Improvements via Linear Regression and Artificial Neural Networks": Supplementary Table S2 Performances of the supplementary linear models using the structural data without sparse atom type pairs.docx

| **Models** | **Number of adopted power exponents in the linear model** | **Scoring power** | | | |
| --- | --- | --- | --- | --- | --- |
|  |  | ***R* in the set composed of 2227 complexes** | ***R* in the subset-1 (1082 complexes from *SET*-1)** | ***R* in the subset-2 (1081 complexes from *SET*-2)** | ***R* in the subset-3 (64 complexes from *SET*-4)** |
| **Supplementary linear model 1** | 1 | 0.652 | 0.656 | 0.634 | 0.817 |
| **Supplementary linear model 2** | 2 | 0.701 | 0.705 | 0.682 | 0.868 |
| **Supplementary linear model 3** | 3 | 0.730 | 0.732 | 0.715 | 0.878 |
| **Supplementary linear model 4** | 4 | 0.764 | 0.772 | 0.744 | 0.905 |
| **Supplementary linear model 5** | 5 | 0.785 | 0.791 | 0.768 | 0.899 |
| **Supplementary linear model 6** | 6 | 0.809 | 0.820 | 0.788 | 0.906 |
| **Supplementary linear model 7** | 7 | 0.830 | 0.838 | 0.814 | 0.913 |
| **Supplementary linear model 8** | 8 | 0.850 | 0.857 | 0.836 | 0.904 |
| **Supplementary linear model 9** | 9 | 0.862 | 0.871 | 0.846 | 0.911 |
| **Supplementary linear model 10** | 10 | **0.877** | **0.888** | **0.862** | **0.887** |
| **Supplementary linear model 11** | 11 | 0.868 | 0.867 | 0.866 | 0.897 |
| **Supplementary linear model 12** | 12 | 0.309 | 0.250 | 0.371 | 0.576 |
| **Supplementary linear model 13** | 13 | 0.238 | 0.194 | 0.297 | 0.602 |
| **Supplementary linear model 14** | 14 | 0.303 | 0.269 | 0.340 | 0.293 |
| **Supplementary linear model 15** | 15 | 0.347 | 0.347 | 0.348 | 0.380 |
| **Supplementary linear model 16** | 16 | 0.294 | 0.286 | 0.296 | 0.294 |
| **Supplementary linear model 17** | 17 | 0.129 | 0.090 | 0.174 | 0.356 |
| **Supplementary linear model 18** | 18 | 0.201 | 0.198 | 0.190 | 0.228 |
