## Supplementary material for "Exploring the Upper Boundaries of Distance-Based Protein–Ligand Binding Affinity Prediction: Accuracy Improvements via Linear Regression and Artificial Neural Networks": Supplementary Table S3 Definition of the heavy atom types and the frequencies of these atom types among the 13733 complexes.docx

| **Atom type** | **Definition** | **Frequency among the complexes** | **Number of involved atom pairs** | **Atom type** | **Definition** | **Frequency among the complexes** | **Number of involved atom pairs** |
| --- | --- | --- | --- | --- | --- | --- | --- |
| C.3 | carbon sp3 | 13733 | 50496267 | S.O2 | sulfone sulfur | 1712 | 418011 |
| C.2 | carbon sp2 | 13733 | 19200985 | P.3 | phosphorus sp3 | 2527 | 1031921 |
| C.1 | carbon sp | 1644 | 360731 | F | fluorine | 1647 | 884608 |
| C.ar | carbon aromatic | 13722 | 35714153 | Cl | chlorine | 1892 | 529116 |
| C.cat | carbocation (C+) used only in a guanidinium group | 11030 | 633347 | Br | bromine | 284 | 106489 |
| N.3 | nitrogen sp3 | 11771 | 1993547 | I | iodine | 144 | 47054 |
| N.2 | nitrogen sp2 | 3651 | 949393 | Fe | iron | 112 | 8302 |
| N.1 | nitrogen sp | 441 | 93735 | Na | sodium | 377 | 5936 |
| N.ar | nitrogen aromatic | 12673 | 5656521 | Mg | magnesium | 755 | 34592 |
| N.am | nitrogen amide | 13733 | 14601264 | K | potassium | 101 | 1432 |
| N.pl3 | nitrogen trigonal planar | 12580 | 3644856 | Ca | calcium | 393 | 12160 |
| N.4 | nitrogen sp3 positively charged | 1769 | 190941 | Mn | manganese | 228 | 9873 |
| O.3 | oxygen sp3 | 13725 | 11758565 | Co.oh | Cobalt (octahedral) | 54 | 2484 |
| O.2 | oxygen sp2 | 13733 | 17198064 | Zn | zinc | 1535 | 42375 |
| O.co2 | oxygen in carboxylate and phosphate groups | 13648 | 4449965 | Se | selenium | 97 | 7875 |
| S.3 | sulfur sp3 | 11917 | 1160858 | B | boron | 228 | 83247 |
| S.2 | sulfur sp2 | 1029 | 267894 | Ir | iridium | 6 | 452 |
| S.O | sulfoxide sulfur | 245 | 10876 | Ni | Nickel | 27 | 312 |
